# RadD from *Fusobacterium nucleatum* Engages NKp46 to Promote Antitumor Cytotoxicity

**DOI:** 10.1101/2025.07.26.666929

**Authors:** Ahmed Rishiq, Johanna Galski, Reem Bsoul, Mingdong Liu, Rema Darawshe, Renate Lux, Gilad Bachrach, Ofer Mandelboim

## Abstract

*Fusobacterium nucleatum*, a Gram-negative bacterium implicated in periodontal disease, has emerged as a contributor to tumor progression in various cancers. Whether the presence of *Fusobacterium nucleatum* inhibits tumor progression is largely unknown. Here, we identify a subspecies-specific interaction between *F. nucleatum* and the natural killer (NK) cell receptor NKp46. Analysis of TCGA datasets revealed that the co-occurrence of *F. nucleatum* and high NKp46 expression correlates with improved survival in head and neck cancers but not in colorectal cancers. Using binding assays, we demonstrate that both human NKp46 and its murine ortholog, Ncr1, directly recognize the fusobacterial adhesin RadD. Genetic deletion of *radD* or blockade of NKp46 significantly impaired NK cell-mediated cytotoxicity *in vitro* and promoted tumor growth. *In vivo* infection with *F. nucleatum* accelerated tumor progression, with an exacerbated effect observed in the absence of RadD or NKp46. These findings highlight RadD as a critical ligand for NKp46 and establish the NKp46–RadD axis as a key interface in host–microbe–tumor interactions, offering a novel target for immunotherapeutic intervention in cancer influenced by microbial factors.

## Introduction

*Fusobacterium nucleatum*, a gram-negative bacterium, has received considerable interest in recent years due to its significance in various human diseases, including cancer (Alon-Maimon et al., 2022). This anaerobic bacterium is mostly found in the human oral cavity (De Andrade et al., 2019). Recent studies showed that *F. nucleatum* influence the progression of various tumor types through its interactions with the host immune system, modulation of inflammatory pathways, and potential involvement in metastatic processes (Guo et al., 2024; Parhi et al., 2020). *F. nucleatum* not only facilitates but also actively promotes breast cancer proliferation and metastasis through established mechanisms — such as invasion of epithelial and endothelial cells via its virulence factors — as well as through pathways that remain to be elucidated (Guo et al., 2024; Parhi et al., 2020). In the context of immune interactions, *F. nucleatum* has been shown to express Fusobacterial apoptosis-inducing protein 2 (Fap2), which engages the TIGIT receptor on natural killer (NK) cells and T cells, leading to immune suppression and inhibition of NK cell cytotoxicity and T cell activity (Gur et al., 2015).

Additionally, another adhesin, CbpF, was found to bind CEACAM1 on T cells modulating their activity (Galaski et al., 2021). We also demonstrated that the RadD protein of *Fusobacterium nucleatum* subsp. *nucleatum* interacts with Siglec-7 on NK cells, leading to the suppression of NK cell-mediated killing of cancer cells (Galaski et al., 2024). In contrast, however, the identity of the *F. nucleatum* ligands that interact with NK activating receptors and how NK cell recognizes this bacterium is still poorly understood. In addition, whether *Fusobacterium nucleatum* could actually inhibits cancer progression of certain tumors is currently unknown.

The natural killer (NK) cell receptor NKp46 has emerged as a focus of research due to its significance in immune response regulation, particularly in the identification and eradication of infected or transformed cells (Barrow et al., 2019). NKp46 was shown to recognize and bind haemagglutinins in both the influenza and the parainfluenza viruses (Mandelboim et al., 2001), Heparan sulfate (HS) and some bacterial and fungal proteins were also identified as ligands for NKp46 (Barrow et al., 2019). Moreover, NKp46 was found to recognize an externalized calreticulin (ecto-CRT), which translocated from the ER to the cell membrane during ER stress (Sen Santara et al., 2023). NKp46 was shown to interact with *F. nucleatum* in the oral cavity (Chaushu et al., 2012). However, the *F. nucleatum* ligand that is recognized by NKp46 and whether the interaction between NKp46 and *F. nucleatum* is important in cancer development and patient’s prognosis remains currently unknown.

Here we demonstrate that the *F. nucleatum* ligand for NKp46 is the RadD adhesin and that this interaction plays a significant role in tumor development.

## Results

### NKp46 Expression Modifies the Prognostic Impact of *F. nucleatum* in a Tumor-Type-Specific Manner

To evaluate the prognostic significance of *Fusobacterium nucleatum* in the context of NKp46 activity, we analyzed transcriptomic data from The Cancer Genome Atlas (TCGA) alongside microbial abundance profiles from The Cancer Microbiome Atlas (TCMA) across two tumor types. In head and neck squamous cell carcinoma (HNSC), patients exhibiting both *F. nucleatum* positivity and high expression of NKp46 (encoded by NCR1) had significantly improved overall survival compared to NKp46⁺ patients lacking *F. nucleatum* (log-rank *P* < 0.05; Figure 1A). Conversely, in colorectal cancer (CRC), *F. nucleatum* status did not significantly stratify survival among NKp46⁺ patients (Figure 1B). The median survival in the *F. nucleatum*⁺NKp46⁺ HNSC subgroup was 5.81 years, compared to 2.36 years in the *F. nucleatum*⁻NKp46⁺ group, corresponding to a hazard ratio (HR) of 2.08 (95% CI: 1.20–3.61; Figure 1C). In CRC, median survival was similar between *F. nucleatum*⁺NKp46⁺ patients (5.15 years) and their *F. nucleatum*⁻ counterparts (5.85 years), with no significant difference in risk (HR = 0.71, 95% CI: 0.26–1.95; Figure 1C). Importantly, NKp46 expression levels were substantially higher in HNSC compared to CRC (Figure 1D), suggesting a possible threshold-dependent effect of NKp46 on microbial-immune interactions. Together, these findings underscore a tumor-type-specific interplay between microbial colonization and immune contexture, positioning NKp46 as a key modulator of *F. nucleatum*–associated clinical outcomes.

**Figure 1:**
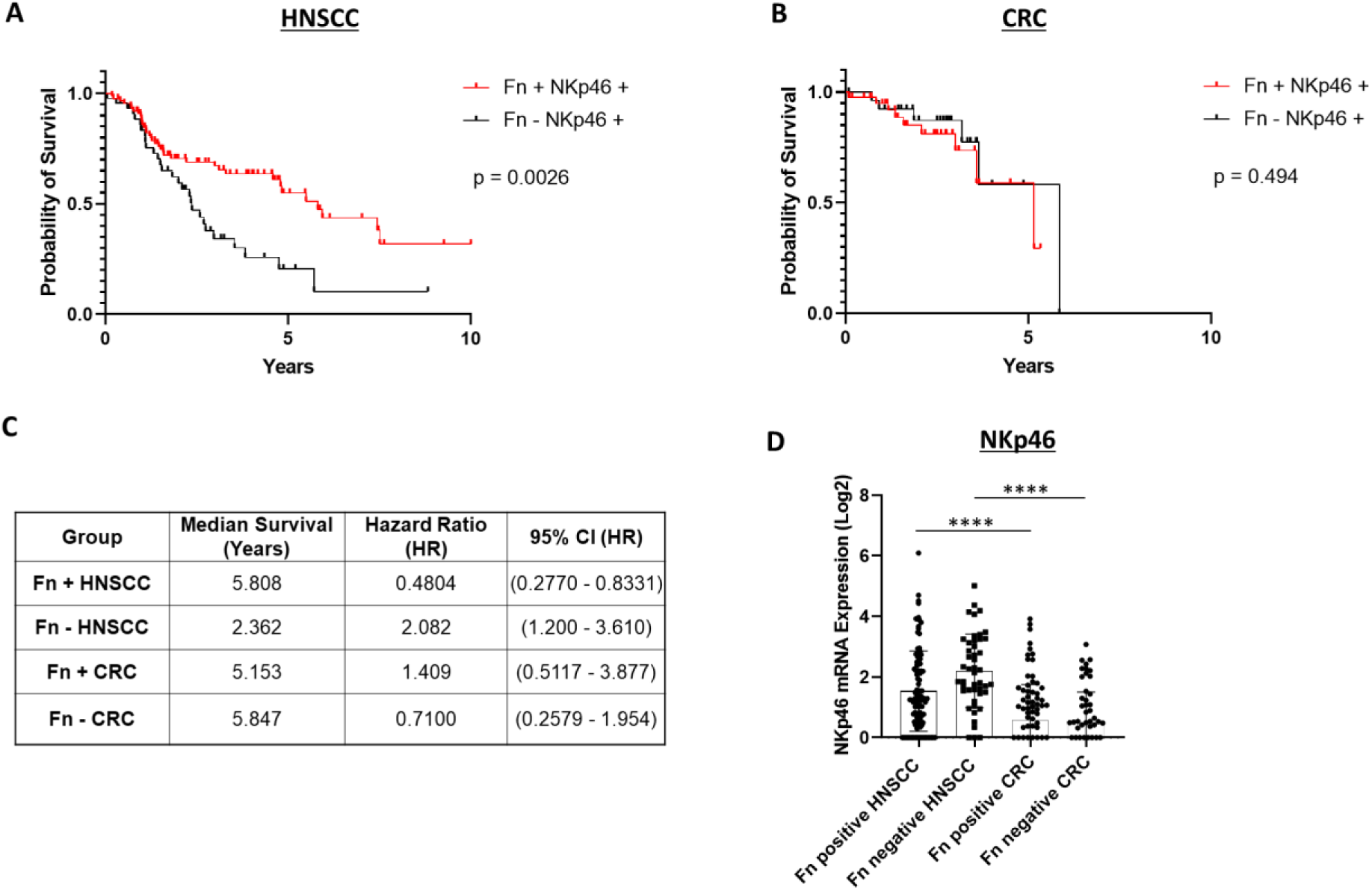
NKp46 expression modifies the prognostic effect of *F. nucleatum* in a tumor-type-specific manner. (A) Kaplan–Meier survival curves for head and neck squamous cell carcinoma (HNSC) patients stratified by *F. nucleatum* status and NKp46 (NCR1) expression. Patients with concurrent *F. nucleatum* positivity (n=87) and high NKp46 expression exhibited significantly improved overall survival compared to those who were *F. nucleatum*-negative (n=44) but NKp46-positive (log-rank *P* < 0.05). (B) Kaplan–Meier survival curves for colorectal cancer (CRC) patients stratified by the same criteria showed no significant difference in survival between *F. nucleatum*–positive (n = 44) and *F. nucleatum*–negative (n = 31) groups. (C) Table summarizing hazard ratios (HR) for *F. nucleatum*–negative cases among NKp46⁺ patients. In HNSC, absence of *F. nucleatum* was associated with a significantly poorer prognosis (HR = 2.08, 95% CI: 1.20–3.61), whereas in CRC, *F. nucleatum* absence showed no significant association with patient prognosis (HR = 0.71, 95% CI: 0.26–1.95). (D) Comparison of NKp46 expression across HNSC and CRC tumors. Log₂ expression levels of NKp46 mRNA were compared across HNSC and CRC cohorts, stratified by *F. nucleatum* positive and negative. Results were analyzed by one-way ANOVA with Bonferroni post hoc correction.

### NKp46 binds RadD

We previously reported that NKp46 interacts with *Fusobacterium nucleatum* and that this interaction plays a role in the context of periodontal disease ^11^. However, the identity of the *F. nucleatum* ligand recognized by NKp46 has remained unknown. Given that co-occurrence of NKp46 expression and *F. nucleatum* presence correlates with improved prognosis in certain tumors—most notably head and neck squamous cell carcinoma—identifying the specific NKp46-binding ligand on *F. nucleatum* is of critical importance for understanding the underlying mechanisms and potential therapeutic implications of this interaction.

To identify the *F. nucleatum* ligand of NKp46, we assessed the binding of NKp46 Ig, its D1 domain (D1 Ig), its mouse orthologue Ncr-1 Ig and CD16 Ig to FITC-labeled *F. nucleatum* strains *ATCC 10953* and *ATCC 23726*, which represent the subspecies *polymorphum* and *nucleatum*, respectively (Figure 2). Surprisingly, NKp46, its D1 domain and the mouse Ncr-1 exhibited higher binding to *ATCC 10953* compared to *ATCC 23726* (Figure 2), while little or no binding was observed for the CD16 which was used as a control.

**Figure 2:**
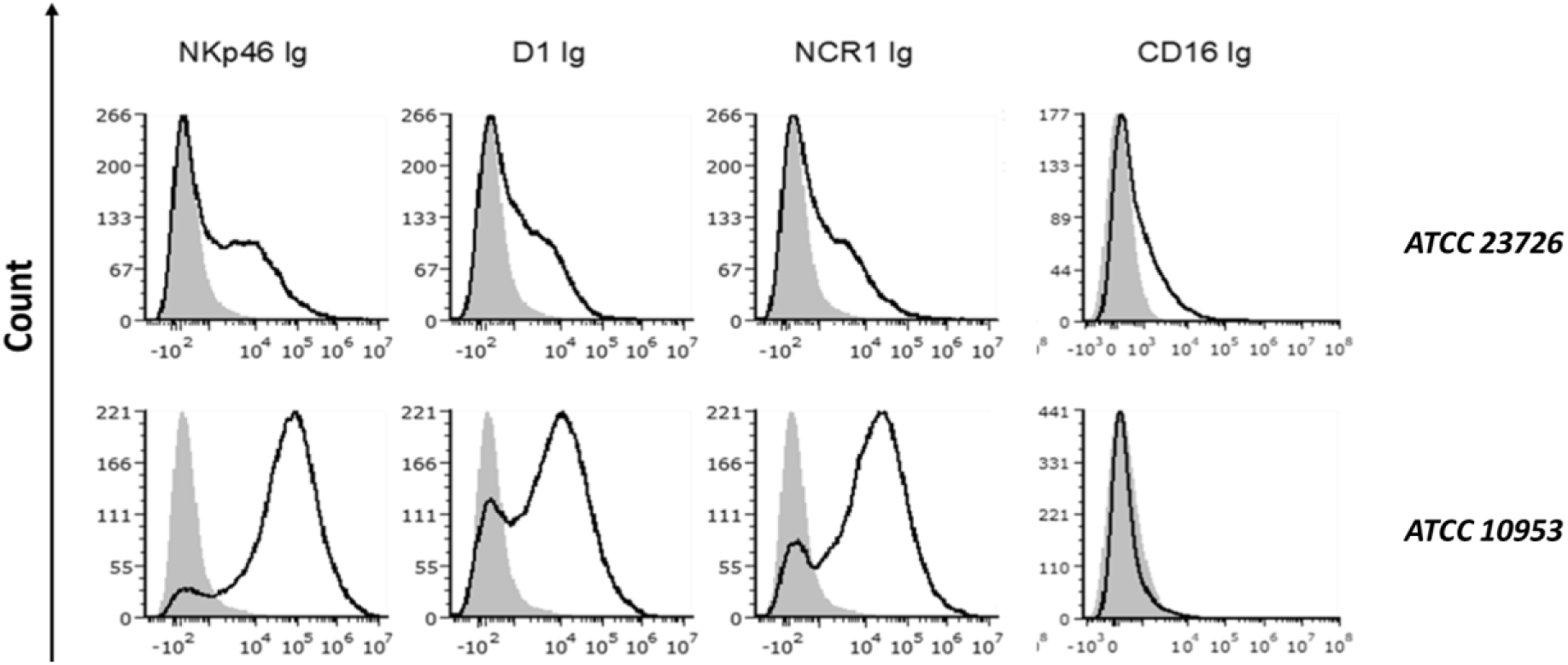
Binding of *Fusobacterium nucleatum* to NKp46 and its D1 domain. The figure shows histograms of FITC-labeled *F. nucleatum subsp. nucleatum ATCC 23726* (FN23726) (upper histograms) and *ATCC 10953* (lower histograms) incubated with 2 μg of NKp46 Ig, D1 domain of NKp46 (D1 Ig), Ncr-1 Ig, and CD16 Ig fusion proteins. Representative staining from one of two independent experiments is shown.

While we were staining various *F. nucleatum* deleted strains with NKp46, we observed an unexpected binding elevation of NKp46 Ig and Ncr1 Ig to FadI deleted mutant (*ATCC 10593 ΔFadI*), while the expression of a control Ig fusion protein CEACAM1 (Ccm-1 Ig) was only minimally increased (please compare Fig 3A and B). Since we showed previously that the absence of FadI results in overexpression of RadD (Figure 3C and (Shokeen et al., 2020)), we incubated Ccm-1 Ig, NKp46 Ig and Ncr-1 Ig with a FITC-labeled *ATCC 10953* mutant lacking the major multifunctional adhesin RadD (*ATCC 10593 ΔRadD*). Interestingly, we observed that the lack of RadD abolished fusobacterial binding of NKp46 Ig and Ncr-1 Ig (Figure 3D).

**Figure 3:**
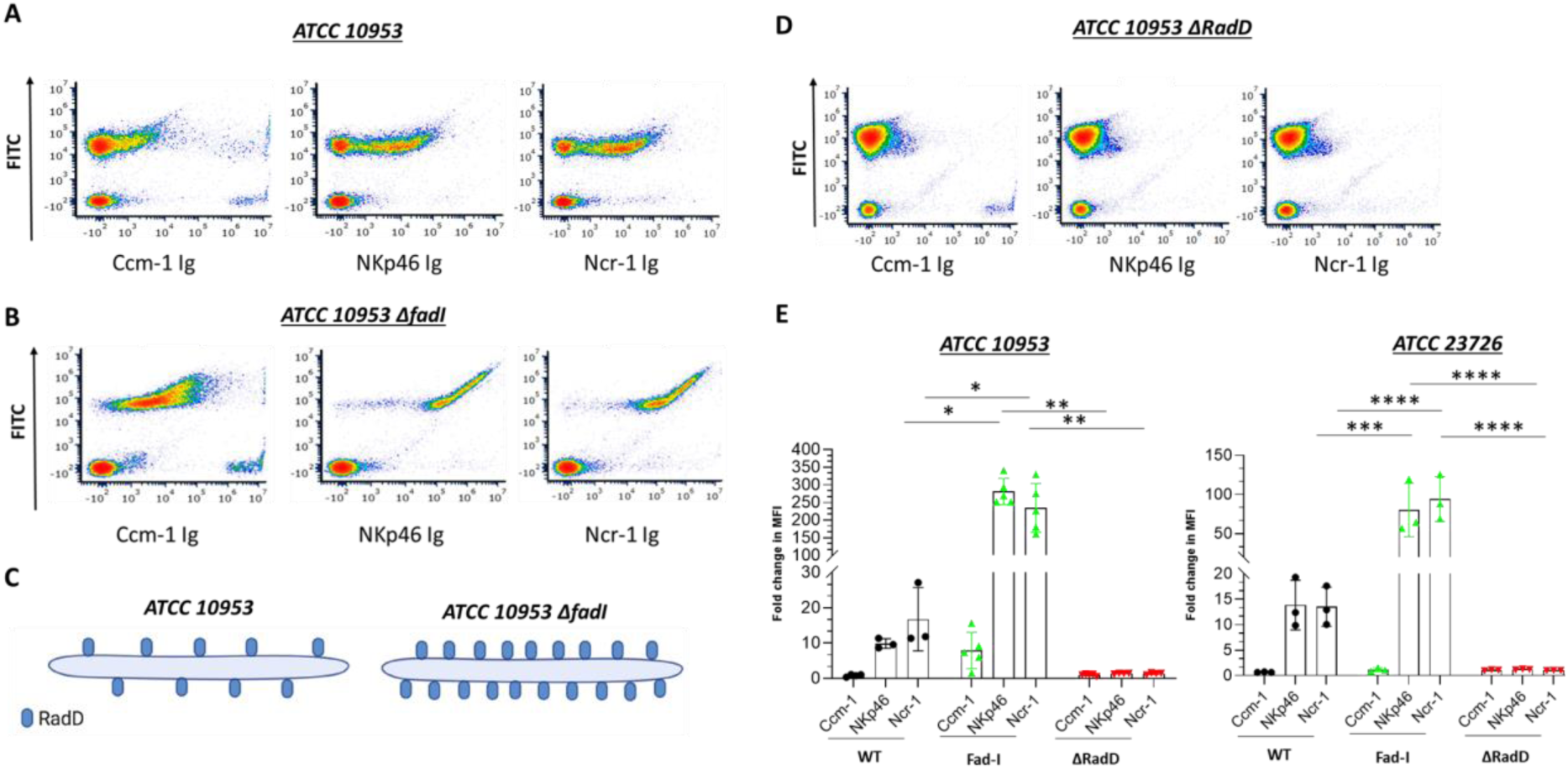
RadD is the bacterial ligand for NKp46. A-B. Density plot of FITC labeled *ATCC 10953* (A) and its *ΔfadI* mutant derivative *ATCC 10953 ΔFad-I* (B) stained with the various fusion proteins (listed in the X axis). C. Schematic representation of *ATCC 10953* wild type (WT) strain and RadD surface expression (Left) compared to *ATCC 10953 ΔFad-I* (Right). D. Density plot of the FITC-labeled *ΔRadD* mutant strain of *ATCC 10953* stained with various fusion proteins (listed in the X axis). The figure shows data from one representative experiment out of three to five independent experiments. E. Fold change quantification of FITC-labeled bacteria binding to the fusion proteins Ccm1-Ig, NKp46 Ig and Ncr-1 Ig in *ATCC 10953* (left) and *ATCC 23726* (right). Summary of three to five independent experiments. The mean value ±SD of the experiments is presented. \**P* < 0.05, \*\**P* ≤ 0.01, \*\*\**P* ≤ 0.001 and \*\*\*\**P* ≤ 0.0001.

Quantification of binding levels confirmed that the absence of FadI resulted in a 250-300-fold increase in binding of NKp46 Ig and Ncr1 Ig to *ATCC 10953 ΔFad-I,* while the absence of RadD almost completely abolished this interaction (Figure 3E, left). A similar effect was observed for the corresponding mutant strains of *ATCC 23726* albeit at a lower level (Figure 3E, right) with *ATCC 23726 Δfad-I* exhibiting an about 100-fold increase, which is abolished in the absence of RadD. These findings indicate that the autotransporter protein RadD is the ligand of NKp46.

### Arginine and anti-NKp46 antibody inhibit the binding of NKp46 to RadD

Since RadD is an arginine-inhibitable protein (Kaplan et al., 2009), we tested whether arginine can block RadD binding to NKp46 or its mouse ortholog Ncr-1. Incubation of FITC-labelled *ATCC 10953* with arginine followed by staining with NKp46 Ig and Ncr-1 Ig revealed that arginine inhibits NKp46 and Ncr-1 Ig binding in a concentration-dependent manner (Figure 4A).

**Figure 4:**
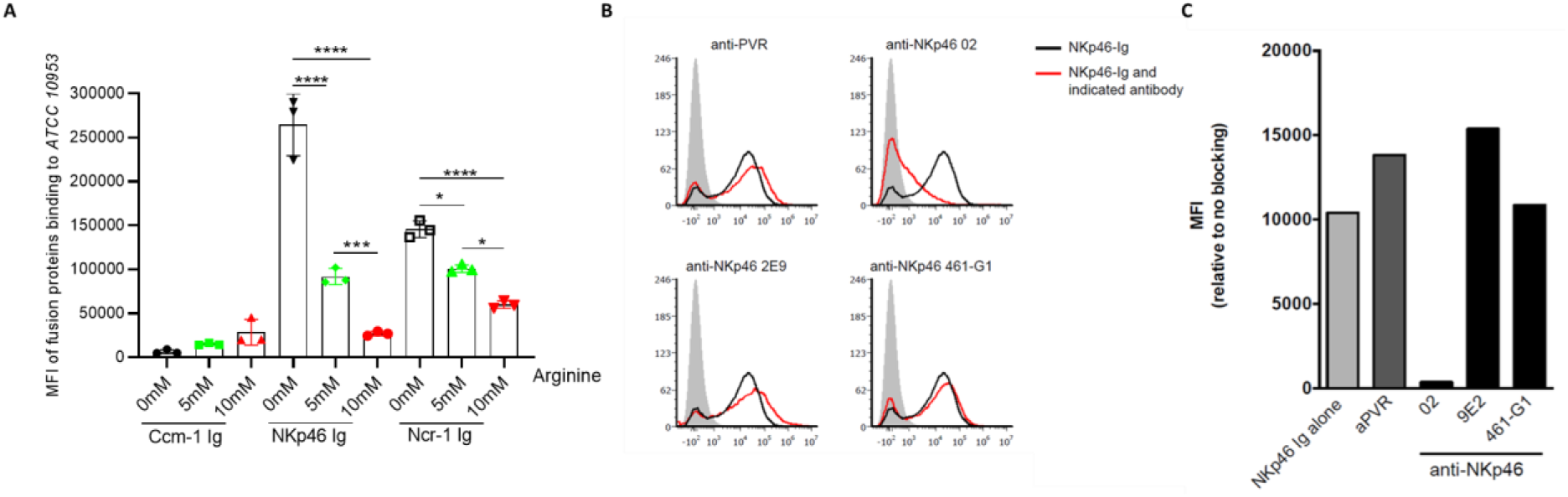
NKp46-02 antibody and arginine block *ATCC 10953* binding to NKp46. A. Quantification of Median Fluorescent Intensity (MFI) of FITC-labeled *ATCC 10953* binding to Ccm1-Ig, NKp46-Ig, and Ncr-1 Ig, without or with 5 and 10 mM of L-Arginine. Data combined from three to four independent experiments are presented. B. NKp46-Ig (2 µg) was pre-incubated with 1 µg of a control anti-PVR antibody and NKp46 monoclonal antibodies (9E2, 461-G1, and 02) to evaluate the blocking of *ATCC 10953* interaction with the NKp46 receptor. C. Shows the quantification results of histograms depicted in A. The mean value ±SD of the experiments is presented. \**P* < 0.05, \*\**P* ≤ 0.01, \*\*\**P* ≤ 0.001 and \*\*\*\**P* ≤ 0.0001.

NKp46 consists of two extracellular domains, a membrane-distal (D1) domain and a membrane-proximal (D2) domain (Barrow et al., 2019). Previous studies indicate that the vast majority of NKp46 ligands are recognized through the domain (D2) (Arnon et al., 2004). Interestingly, *F. nucleatum* seemed to be recognized by the D1 domain of the NKp46 receptor (Figure 2). To test whether the D1 domain of NKp46 is involved in its binding to *F. nucleatum* we used several anti-human NKp46 antibodies (461-G1, hNKp46.02 (02), and 9E2) that were previously shown to bind the D1 domain of NKp46 (Berhani et al., 2019). We pre-incubated NKp46 Ig individually with all of these antibodies prior to addition to *ATCC 10953*. Notably, no difference in binding of *ATCC 10953* was observed when the NKp46 Ig was incubated with either 9E2, 461-G1, or anti-PVR antibody which served as controls (Figure 4B). However, blocking with hNKp46.02 (02) antibody significantly reduced the NKp46 Ig-*ATTC 10953* interaction (Figure 4B, quantified in 4C). These findings confirm that the NKp46 receptor interacts with *ATCC 10953* specifically via its D1 domain.

### NKp46-RadD interactions lead to tumor cell killing *in vitro* and *in vivo*

Next, we examined the impact of the NKp46.02 (02)-blocking antibody on NK cell cytotoxicity against tumor cells incubated with *F. nucleatum*. Hence, we co-incubated or not NK cells with the 02 antibody. Subsequently, NK cells were co-cultured with human mammary gland carcinoma cell lines (MCF7, T47D) that were incubated with *ATCC 10953* or with the corresponding *ΔRadD* mutant strains. NK cytotoxicity was assessed using the Calcein-AM assay (Figure 5A). We observed that the 02 antibody had no effect on NK cell cytotoxicity against breast cancer cell lines, in the absence of *F. nucleatum*. However, a significant increase in tumor cell killing (approx.1.2-1.5-fold change) was noticed for unblocked NK cells compared to 02 blocked NK cells (Figure 5B and 5C). Interestingly, this increased NK cytotoxicity was diminished when the tumor cells were incubated with *ATCC 10953 ΔRadD* and in the absence of RadD NKp46 blocking had no effect (Figure 5B and 5C).

**Figure 5:**
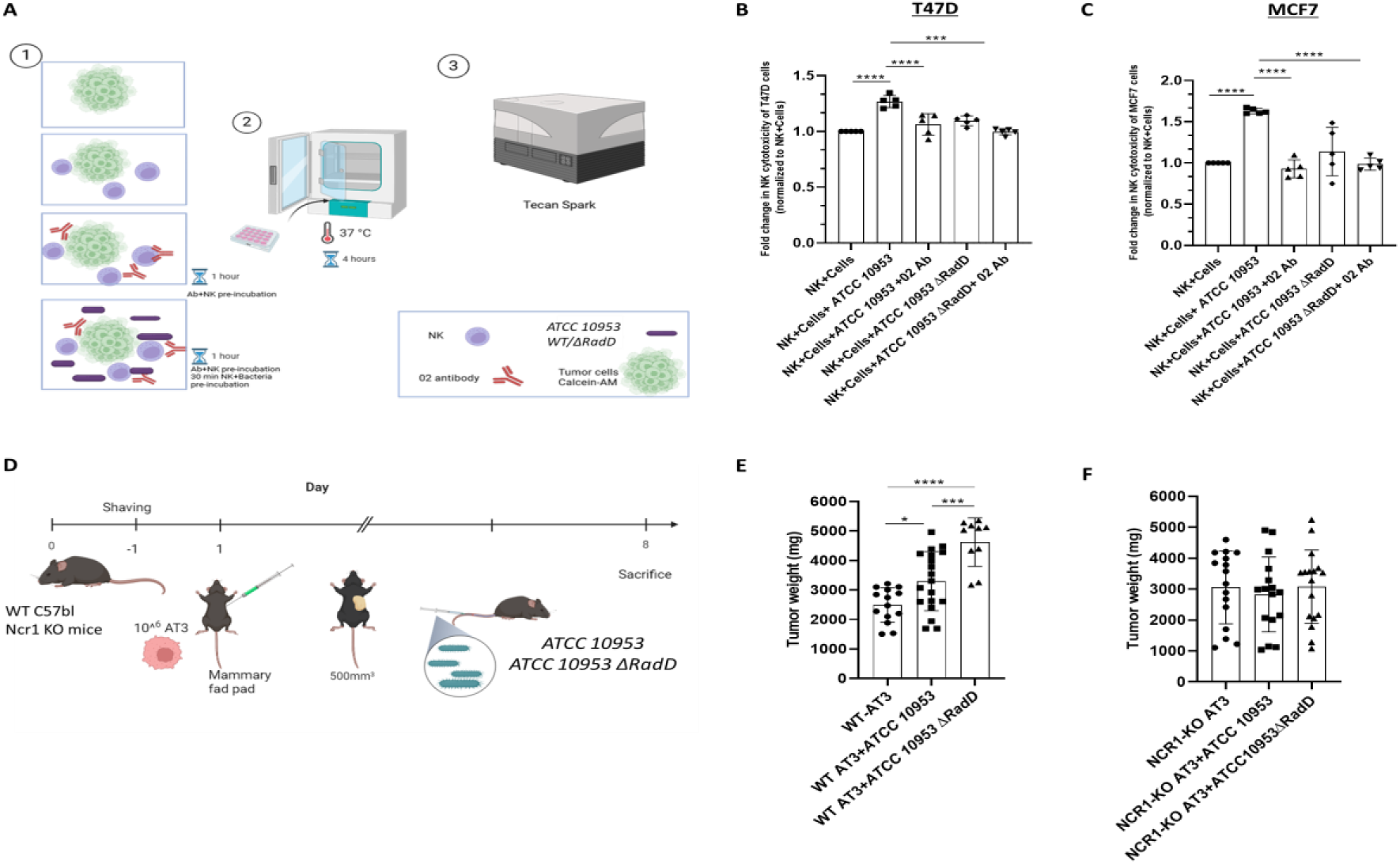
Cytotoxicity and tumor growth is RadD and Ncr1-dependent. A. Schematic diagram showing the design of the NK cells cytotoxicity assay against breast cancer cell lines T47D and MCF7. 1. Tumor cells were stained with Calcein-AM dye and then incubated either with tumor cells (T47D or MCF7) only, tumor + NK, tumor + bacteria (*ATCC 10953*WT and *ATCC 10953 ΔRadD*) + NK with/without preincubation with 02 antibody. 2. Killing assays were performed in a 37°C incubator for 4 hours. 3. The fluorescence intensity of Calcein was measured to determine cell viability using a spectrophotometer (Tecan Spark). Summary of NK cytotoxicity against T47D (B) and MCF7 (C) breast cancer cell lines. Combined results from five independent experiments. D. C57BL\6 or NCR1-KO mice were shaved and AT3 cells (1 x 10^^6^ cells in 100 μl PBS) were injected one day later into the mammary fat pad. When tumors reached a size of about 500 mm^3^, mice were inoculated intravenously with 5 × 10^7^ *ATCC 10953*WT and 7.5 × 10^7^ *ATCC 10953* Δ*RadD* bacteria. Eight days later, mice were sacrificed and tumor weight was determined. E. The tumor weight of C57BL or NCR1-KO (F) mice. The figure shows the combination of 4-5 experiments performed. The mean value ±SD of the experiments is presented. \**P* < 0.05, \*\**P* ≤ 0.01, \*\*\**P* ≤ 0.001 and \*\*\*\**P* ≤ 0.0001.

To test whether the NKp46-RadD interactions are important for controlling tumor growth, we established a syngeneic mouse breast cancer model by implanting AT3 cell line orthotopically in the mammary fat pad of C57BL\6 wild type (WT) and Ncr-1 deficient mice (NCR-1 KO). The tumor was allowed to grow to reach approximately 500 mm^3^ in volume prior to intravenous inoculation with either *ATCC 10953* or *ATCC 10953 ΔRadD*. Mice were sacrificed on day 8 following the bacteria injection and tumor weight was measured (Figure 5D). The tumor weight was significantly increased when wild type *ATCC 10953* was inoculated as compared to uninfected tumor-bearing WT mice (Figure 5E). Strikingly, a further increase in tumor weight was observed when mice were injected with *ATCC 10953 ΔRadD* mutant bacterium (Figure 5E). The increased tumor growth was not observed in the NCR-1 KO mice (Figure 5F). These results indicate that *RadD* recognition by the NKp46 activating receptor is required for better cytotoxicity against tumors infected with *F. nucleatum*.

## Discussion

Patient prognosis is a key determinant in the association between *Fusobacterium nucleatum* infection and tumour development and progression, for instance, in colorectal cancer (CRC) metastasis (Lee et al., 2021). To date, no evidence has clarified whether NKp46 activation in response to *F. nucleatum* plays a significant role in influencing patient survival. Our analysis of TCGA PanCancer and TCMA datasets revealed that co-occurrence of *F. nucleatum* and NKp46 expression is associated with a protective effect in head and neck squamous cell carcinoma (HNSC) patients. In contrast, no such association was observed in colorectal cancer (CRC) cohorts, likely due to markedly lower NKp46 expression levels in these patients.(Cerami et al., 2012; de Bruijn et al., 2023; Dohlman et al., 2021; Gao et al., 2013).

In this study, we demonstrate that the RadD protein of *F. nucleatum* is specifically recognized by the D1 domain of the NKp46 receptor. This interaction enhances the ability of NK cells to kill tumor cells infected with *F. nucleatum*. Previous studies have shown that *F. nucleatum* promotes tumor growth in both breast (Parhi et al., 2020) and colorectal cancers (Zhu et al., 2024), and suppresses immune cell activity through engagement of three inhibitory receptors: CEACAM1, TIGIT, and Siglec-7 (Galaski et al., 2024, 2021; Gur et al., 2015). Furthermore, it was also shown that CD147 which is overexpressed on the surface of colorectal cancer (CRC) cells also binds to RadD. The binding of RadD to CD147 leads to the enrichment of *F. nucleatum* within CRC tissues, triggering an oncogenic cascade PI3K– AKT–NF-κB signaling pathway, which promotes tumorigenesis (Zhang et al., 2024). Here, we uncover a counterbalancing mechanism, whereby NK cells overcome this immune suppression through NKp46-mediated recognition of RadD and that in the absence of this interaction tumor growth is even further increased.

*F. nucleatum* RadD is an outer membrane autotransporter protein that mediates binding between *F. nucleatum* and other oral bacteria by promoting interspecies interactions, which facilitates dental plaque development and virulence in periodontal disease (Kaplan et al., 2009). RadD in combination with Fap2, have been identified as virulence factors capable of inducing cell death in lymphocytes (Kaplan et al., 2010). As an adhesin, RadD facilitates the coaggregation of *F. nucleatum* synergistically with *Clostridioides difficile* (*C. difficile*). This interaction promotes biofilm formation within the intestinal mucus, potentially contributing to the pathogenesis of *C. difficile* infection (Engevik et al., 2021). Fusobacterium-associated defensin inducer (Fad-I) is a cell wall-associated diacylated lipoprotein from *F. nucleatum* that acts as a key microbial molecule that enhances the host’s innate immune response at mucosal surfaces by promoting human beta-defensin (hBD-2) expression (Bhattacharyya et al., 2016). Inactivation of the *fad-I* (*Δfad-I*) gene results in a significant increase in *radD* gene expression, leading to elevated *radD* transcript levels and subsequently the binding to *Streptococcus gordonii* was increased (Shokeen et al., 2020).

While NKp46 showed strong binding to *F. nucleatum* subsp. *polymorphum (ATCC 10953)*, less binding was observed for *F. nucleatum* subsp. *nucleatum (ATCC 23726)*. The differences in binding could be due to substantial differences in RadD expression between the different *F. nucleatum* subspecies.

Given that deletion of *F. nucleatum fad-I* resulted in elevated surface expression of RadD (Shokeen et al., 2020), we noticed enhanced binding to both human and mouse NKp46 receptors. However, this interaction was abolished in the *F. nucleatum ΔRadD* strain, identifying RadD as NKp46 ligand. Consistent with previous studies, we revealed that NKp46 binding to RadD is arginine-dependent (Kaplan et al., 2009), which not only impairs *F. nucleatum* adhesion and biofilm formation but also reduces its interaction with NKp46.

Importantly, using the NKp46 blocking antibody (02 mAb), which targets the D1 domain of the receptor (Berhani et al., 2019), we demonstrated that this antibody effectively disrupts *F. nucleatum* binding to NKp46 and impairs NK cell-mediated killing against tumor cells. In our *in vivo* model, WT mice infected with the *ATCC 10953ΔRadD* strain exhibited significantly greater tumor weight relative to those infected with the wild-type *ATCC 10953* strain.

Intriguingly, infection with either the wild-type *ATCC 10953* or *ATCC 10953* Δ*RadD* strains was not able to affect tumor progression in NCR-1 KO mice. These results collectively support our hypothesis that NK cell cytotoxicity is mediated by the presence of NKp46 on NK cells and the expression of RadD on the *F. nucleatum* surface (Figure 6).

**Figure 6:**
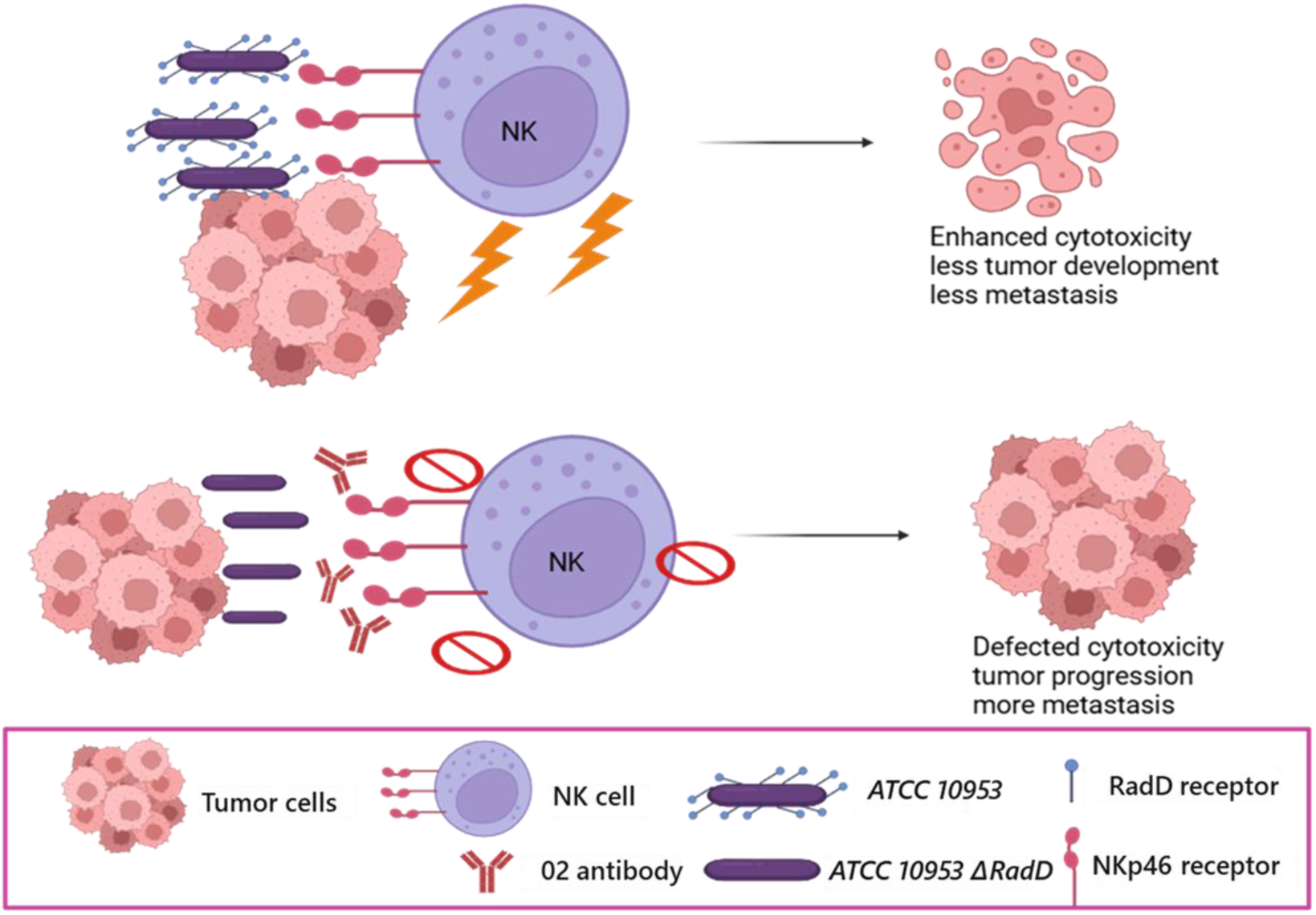
Postulated model for the RadD-NKp46 interaction impact on NK cytotoxicity and tumor growth. NKp46 interaction with RadD expressed by *Fusobacterium nucleatum* triggers NK cell cytotoxicity. This activation enhances tumor cell killing *in vitro* and *in vivo*. Conversely, the absence of RadD or the blocking of NKp46 impairs NK cell activity, leading to tumor progression. Created by BioRender.

Our findings align with those of another group that investigated the *ΔRadD* mutant in a mouse model of preterm birth (Wu et al., 2021). Using the *ATCC 23726* strain the authors revealed earlier and increased invasion of the *ΔRadD* strain which reached the placenta, amniotic fluid, and fetus sooner and continued accumulating over time in comparison with the WT *F. nucleatum.* Moreover, the RadD mutant exhibited reduced systemic clearance, with no decline observed in liver or spleen levels, suggesting impaired immune evasion or a lack of control over dissemination (Wu et al., 2021).

In conclusion, we demonstrate that *Fusobacterium nucleatum* RadD functions as a direct ligand for NKp46 and highlight its critical role in modulating NK cell activity both *in vitro* and *in vivo*. Strategies aimed at elevating NKp46 expression or enhancing its activity might further strengthen NK cell responses and help reduce cancer development associated with *F. nucleatum* infection.

## Materials and Methods

### Key resources table

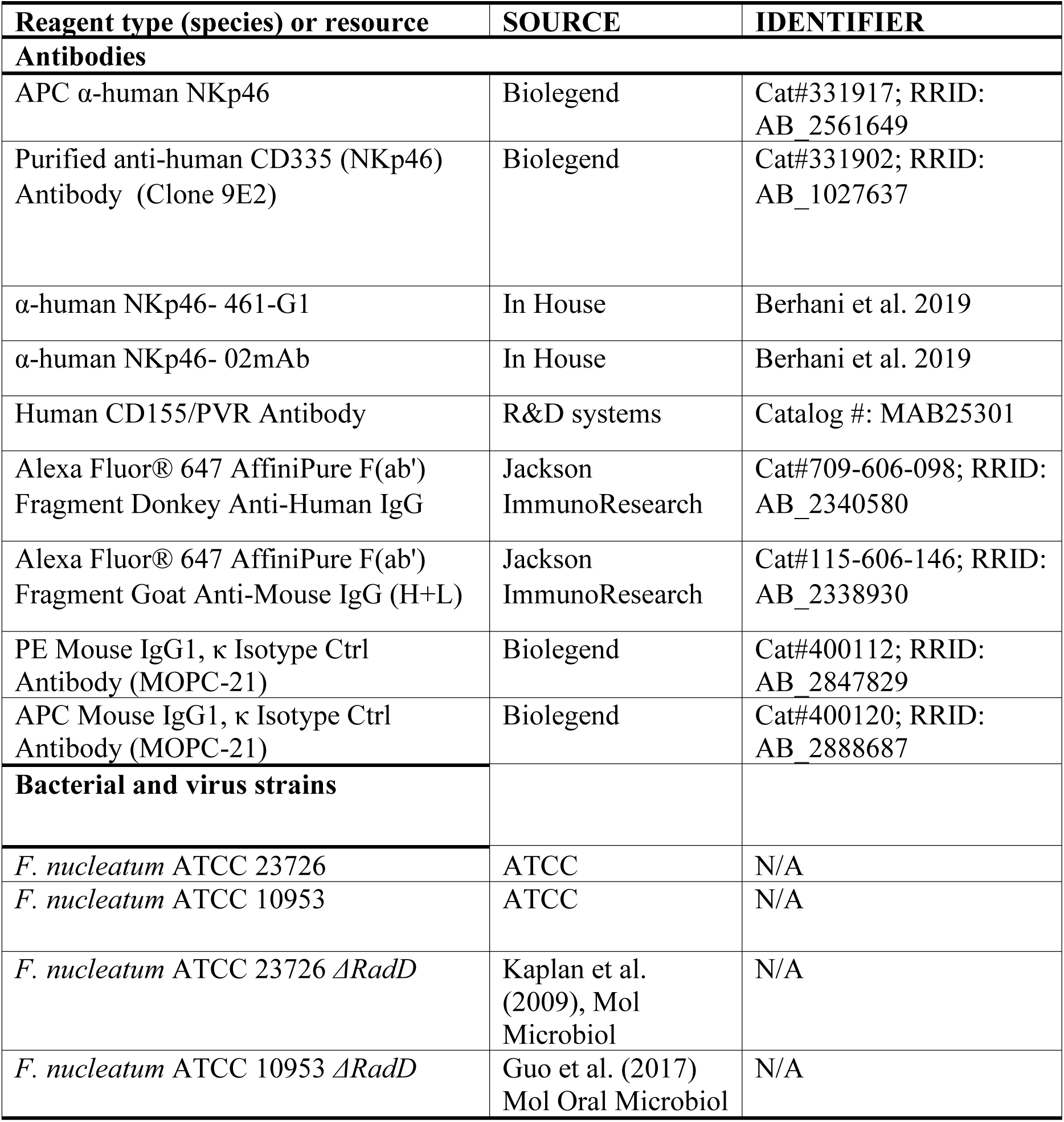
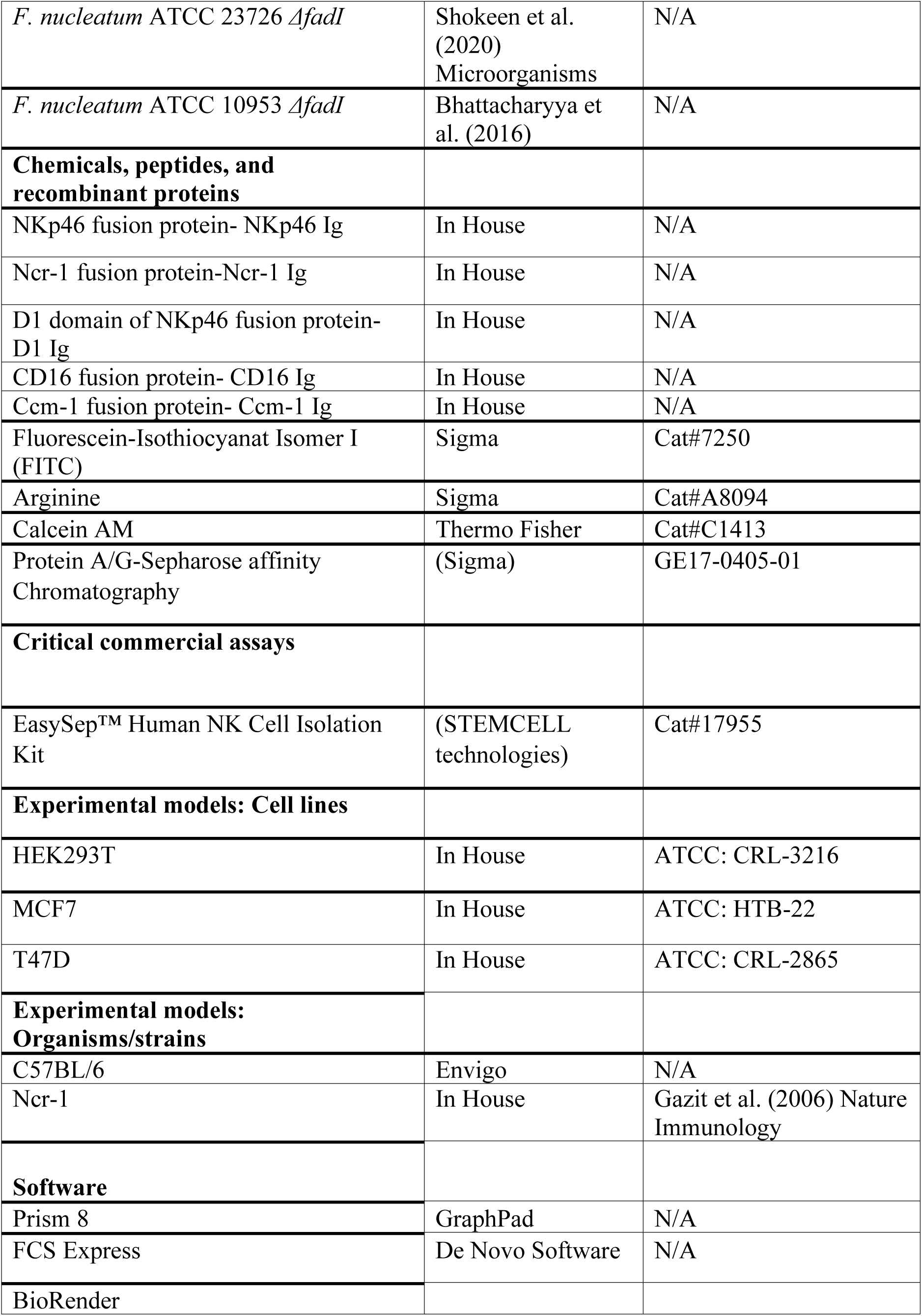

### TCGA and TCMA

RNA-sequencing expression data and corresponding clinical and survival information were retrieved from the PanCancer Atlas dataset available through cBioPortal (https://www.cbioportal.org/). Data for head and neck squamous cell carcinoma (HNSC) and colorectal cancer (CRC) were selected for analysis. Microbial abundance scores for *Fusobacterium nucleatum* were curated from The Cancer Microbiome Atlas (TCMA). Co-occurrence of NKp46 expression and *F. nucleatum* abundance was evaluated using survival analyses (Kaplan–Meier with log-rank test using Prism were Statistical significance was defined as *P* < 0.05.

### Cell lines

C57BL\6 mouse mammary carcinoma cell line (AT3), human breast mammary gland adenocarcinoma MCF7, and T47D cell lines were cultured with DMEM or RPMI with 10% inactivated fetal bovine serum (Sigma-Aldrich), 1 mM sodium pyruvate (Biological Industries), 2 mM glutamine (Biological Industries), nonessential amino acids (Biological Industries), 100 U/ml penicillin (Biological Industries) and 0.1 mg/ml streptomycin (Biological Industries). All cells were tested regularly for mycoplasma using Mycolor One-Step Mycoplasma Detector (Vazyme) kit. Primary NK cells were isolated from the human peripheral blood of healthy individuals using EasySep™ Human NK Cell Isolation Kit (STEMCELL technologies) and then cultured in F12-DMEM medium supplemented with 10% human serum (Sigma-Aldrich), 1 mM sodium pyruvate (Biological Industries), 2 mM glutamine (Biological Industries), nonessential amino acids (Biological Industries), 100 U/ml penicillin (Biological Industries), 0.1 mg/ml streptomycin (Biological Industries) and 400IU of recombinant human hIL2 (Peprotech).

### Fusion proteins

To generate fusion proteins, the extracellular portion of the protein of interest was cloned into a mammalian expression vector containing the mutated Fc portion of human IgG1 (CSI-Ig IRES-Puro Fc mut N197A). Fusion proteins NKp46 Ig, CD16-Ig, D1-Ig, Ncr-1 Ig were generated in HEK293T cells and purified using Protein A/G-Sepharose affinity Chromatography (Sigma-Aldrich).

### Bacteria cultivation

The bacterial strains used in this study were *F. nucleatum* subsp. *nucleatum ATCC 23726*, *F. nucleatum* subsp. *polymorphum ATCC 10953* and their respective *ΔRadD* and *ΔfadI* mutant derivatives ^21^. Bacteria were kept in −80°C frozen glycerol stocks and grown at 37°C on blood agar plates (Hylabs) under anaerobic conditions generated using the Oxoid AnaeroGen anaerobic gas generator system (Thermo Fisher). Bacteria were harvested from blood agar plates for subsequent experimental procedures.

### Bacteria staining and flow cytometry

In brief, for bacteria staining experiments bacteria were harvested from blood agar plates, washed twice with PBS (Sartorius), and incubated with 0.1 mg/ml FITC (Sigma-Aldrich) in PBS at room temperature in the dark for 30 minutes on a shaker. Subsequently, bacteria were washed thrice in PBS at 4,000 rpm for 10 minutes to remove unbound FITC. Next, bacteria were divided into 96-well U plates at 2 million bacteria per well and incubated with 2 μg of fusion proteins per well for 1 hour on ice, followed by washing and 30 minutes of incubation with Alexa Fluor 647-conjugated donkey anti-human IgG (Jackson Immunoresearch). Histograms of bacteria were gated on FITC-positive cells.

For arginine-blocking experiments, incubation of *ATCC 10953* was performed in the presence of arginine (5 mM or 10 mM) for 30 minutes. Subsequently, 2ug of Ccm-1 Ig, NKp46 Ig, or Ncr-1Ig were added for another 30 minutes. Bacteria were centrifuged at 4,000 rpm for 10 minutes, washed with PBS, and stained with Alexa Fluor 647-conjugated donkey anti-human IgG (Jackson Immunoresearch) for 30 minutes on ice.

Antibody-blocking experiments were performed by incubating 2 μg of NKp46 Ig with 1 μg of α-human NKp46 (clones 9E2, 461-G1, and 02) or anti-PVR antibodies for 1 hour on ice. Subsequently, this incubation was followed by washing 2 times with PBS and 30 minutes with Alexa Fluor 647-conjugated donkey anti-human IgG.

### In vitro cytotoxicity

NK cytotoxicity was investigated as previously described (Galaski et al., 2024). We found that for *ATCC 10953 ΔRadD* an MOI 25:1 was necessary to achieve adhesion to the breast cancer cell line that was similar to MOI 10:1 for the control *ATCC 10953* strain. Effector NK cells (100,000 cells) isolated from healthy individuals was incubated with or without 2 ug of anti-NKp46 (02) antibody in antibiotic-free RPMI medium for 1 hour on ice. Then, *ATCC 10953* and *ATCC 10953 ΔRadD* were incubated with NK cells for 30 minutes in a 37°C incubator. Calcein-AM (Thermoscientific) stained tumor cell lines were cocultured with the effector NK cells in a 10:1 effector-to-target ratio for an additional 4 hours in a 37°C incubator. The maximal killing was determined by adding Triton-X (9.5 ml RPMI+ 0.5 ml Triton-X) to the target cells (with or without bacteria), and the spontaneous release was determined by adding only target cells (with or without bacteria). Plates were centrifuged (1,600 rpm for 5 min, 4°C), and supernatants (75 μl) were transferred to a black 96-well plate. The Calcein-AM release into the supernatant was measured using a Tecan Spark multiplate reader with excitation/emission wavelengths at 485nm/535 nm. Specific lysis percentage was calculated as follows: 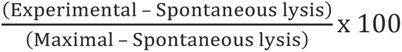. The fold change was calculated by normalizing the experimental groups to the NK and tumor cells (with or without 02 antibody).

### In vivo experiments

7-8 weeks-old female wild-type C57BL/6 and Ncr-1 knockout mice (NCR-1 KO) were injected orthotopically (mammary fat pad) with 1 × 10^6^ AT3 tumor cells. At a tumor size of 500 mm^3^, mice were randomly divided into three groups and injected intravenously with 5 × 10^7^ *F. nucleatum* ATCC 10953, 7.5 × 10^7^ *F. nucleatum* ATCC 10953 *ΔRadD*, and one group of AT3 tumor cells only. Mice were sacrificed at day 8, and tumor weight was measured.

### Statistical analysis

Statistical analysis and graphs were prepared using GraphPad Prism version 8 (GraphPad Software). Comparison among groups was performed by one-way ANOVA multiple comparison test followed by Tukey’s post-hoc test. *P*-value was considered significant at *P* < 0.05.

## Acknowledgements

This work was supported by the following grants awarded to O.M.: the ICRF professorship grant, the ISF grant 619/23, the Israeli Innovation Authority grants 72670 and 75934.

## Author contributions

A.R., J.G. and O.M. conceived the study. A.R., J.G., M.L., R.B., and R.D. performed the experiments. A.R., J.G., R.B. and O.M. analyzed the data. R.L. contributed *F. nucleatum* mutants and analyses of *Fusobacterium* genomic information of autotransporter-like large outer membrane proteins. A.R. drafted the manuscript. O.M. and G.B. supervised the project. All authors revised and approved the manuscript.

